# Evolution of metastases-associated fibroblasts in the lung microenvironment is driven by stage-specific transcriptional plasticity

**DOI:** 10.1101/778936

**Authors:** Ophir Shani, Yael Raz, Lea Monteran, Or Megides, Hila Shacham, Noam Cohen, Dana Silverbush, Oshrat Levi-Galibov, Camilla Avivi, Roded Sharan, Asaf Madi, Ruth Scherz-Shouval, Iris Barshack, Ilan Tsarfaty, Neta Erez

**Affiliations:** Department of Pathology, Sackler Faculty of Medicine, Tel Aviv University, Tel Aviv, Israel; Department of Obstetrics and Gynecology, Tel Aviv Sourasky Medical Center, Tel Aviv, Israel; Department of Clinical Microbiology and Immunology, Sackler Faculty of Medicine, Tel Aviv University, Tel Aviv, Israel; Blavatnik School of Computer Sciences, Faculty of Exact Sciences, Tel Aviv University, Tel Aviv, Israel; Department of Biomolecular Sciences, The Weizmann Institute of Science, Rehovot, Israel; Department of Pathology, Sheba Medical Center, Tel Hashomer, affiliated with Sackler Faculty of Medicine, Tel Aviv University, Tel Aviv, Israel

**Keywords:** Brest cancer, metastasis, fibroblasts, microenvironment

## Abstract

Mortality from breast cancer is almost exclusively a result of tumor metastasis, and lungs are one of the main metastatic sites. Cancer-associated fibroblasts (CAFs) are prominent players in the microenvironment of breast cancer. However, their role in the metastatic niche is largely unknown. In this study, we profiled the transcriptional co-evolution of lung fibroblasts isolated from transgenic mice at defined stage-specific time points of metastases formation. Employing multiple knowledge-based platforms of data analysis provided powerful insights on functional and temporal regulation of the transcriptome of fibroblasts. We demonstrate that fibroblasts in lung metastases are transcriptionally dynamic and plastic, and reveal stage-specific gene signatures that imply functional tasks, including extracellular matrix remodeling, stress response and shaping the inflammatory microenvironment. Furthermore, we identified *Myc* as a central regulator of fibroblast rewiring and found that stromal upregulation of *Myc* transcriptional networks is associated with worse survival in human breast cancer.

## Introduction

Breast cancer continues to be one of the leading causes of cancer related death in women, and mortality is almost exclusively a result of tumor metastasis. Advanced metastatic cancers are mostly incurable and available therapies generally prolong life to a limited extent. It is increasingly appreciated that in addition to tumor cell-intrinsic survival and growth programs, the microenvironment is crucial in supporting metastases formation ^1-3^. Nevertheless, while years of research have revealed the central role of the microenvironment in supporting tumor growth and response to therapy at the primary tumor site ^3-5^, the role of the metastatic microenvironment and the molecular crosstalk between stromal cells, including fibroblasts and immune cells at the metastatic niche are poorly characterized.

Preparation of secondary sites to facilitate subsequent tumor cell colonization has been described for multiple cancers ^6^. Secreted factors and extracellular vesicles from tumor and stromal cells were reported to instigate a permissive pre-metastatic niche by influencing the recruitment and activation of immune cells ^7-11^, and by modifying the composition of the extracellular matrix (ECM) ^12-16^. Each metastatic microenvironment exerts specific functions that support or oppose colonization by disseminated tumor cells ^6,17^. Therefore, understanding distinct organ-specific mechanisms that enable metastatic growth is of crucial importance.

Lungs are one of the most common sites of breast cancer metastasis. Various immune cell populations were shown to be functionally important in facilitating breast cancer pulmonary metastasis ^10,18-21^. However, very little is known about the role of fibroblasts during the complex process of metastases formation.

Cancer-associated fibroblasts (CAFs) are a heterogeneous population of fibroblastic cells found in the microenvironment of solid tumors. In some cancer types, including breast carcinomas, CAFs are the most prominent stromal cell type, and their abundance correlates with worse prognosis ^22^. We previously demonstrated a novel role for CAFs in mediating tumor-promoting inflammation in mouse and human carcinomas ^23,24^. We further characterized the origin, heterogeneity and function of CAFs in breast cancer ^25-27^. Importantly, we found profound changes in the expression of pro-inflammatory genes in fibroblasts isolated from metastases-bearing lungs ^26^. However, comprehensive profiling of metastases-associated fibroblasts was not previously done. Based on the central role of CAFs in supporting tumor growth at the primary tumor site ^28^, we hypothesized that transcriptional reprogramming of lung fibroblasts is an important factor in the formation of a hospitable metastatic niche that supports breast cancer metastasis.

In this study, we set out to characterize the dynamic co-evolution of fibroblasts during pulmonary metastasis. To achieve this goal, we utilized novel transgenic mice that enable visualization, tracking, and unbiased isolation of fibroblasts from spontaneous lung metastases. Here we demonstrate the profiling and analysis of the dynamic evolution of fibroblast transcriptome at distinct disease stages, including early and late metastatic disease.

## Results

### Fibroblasts are activated and transcriptionally reprogrammed in the lung metastatic niche

We previously demonstrated that fibroblasts at the primary tumor microenvironment are reprogrammed to obtain a pro-inflammatory and tumor-promoting phenotype ^24,25,27^. Moreover, we found that fibroblasts are also modified at the lung metastatic niche ^26^. In this study, we set out to characterize the changes in lung fibroblasts that mediate the formation of a hospitable niche in breast cancer metastasis.

We initially investigated metastases-associated fibroblasts in the lung metastatic microenvironment of MMTV-PyMT transgenic mice with spontaneous lung metastases, compared with normal lungs. We analyzed the changes in the population of fibroblasts using immunostaining with multiple known fibroblast markers including αSMA, FSP-1^29,30^ and Podoplanin (PDPN) ^31^ (Figure 1A-D). Notably, analysis of αSMA and FSP-1 indicated an upregulation in the expression of these markers in metastases-bearing lungs (Figure 1B,C), suggesting that lung metastases are associated with fibroblast activation.

**Figure 1.**
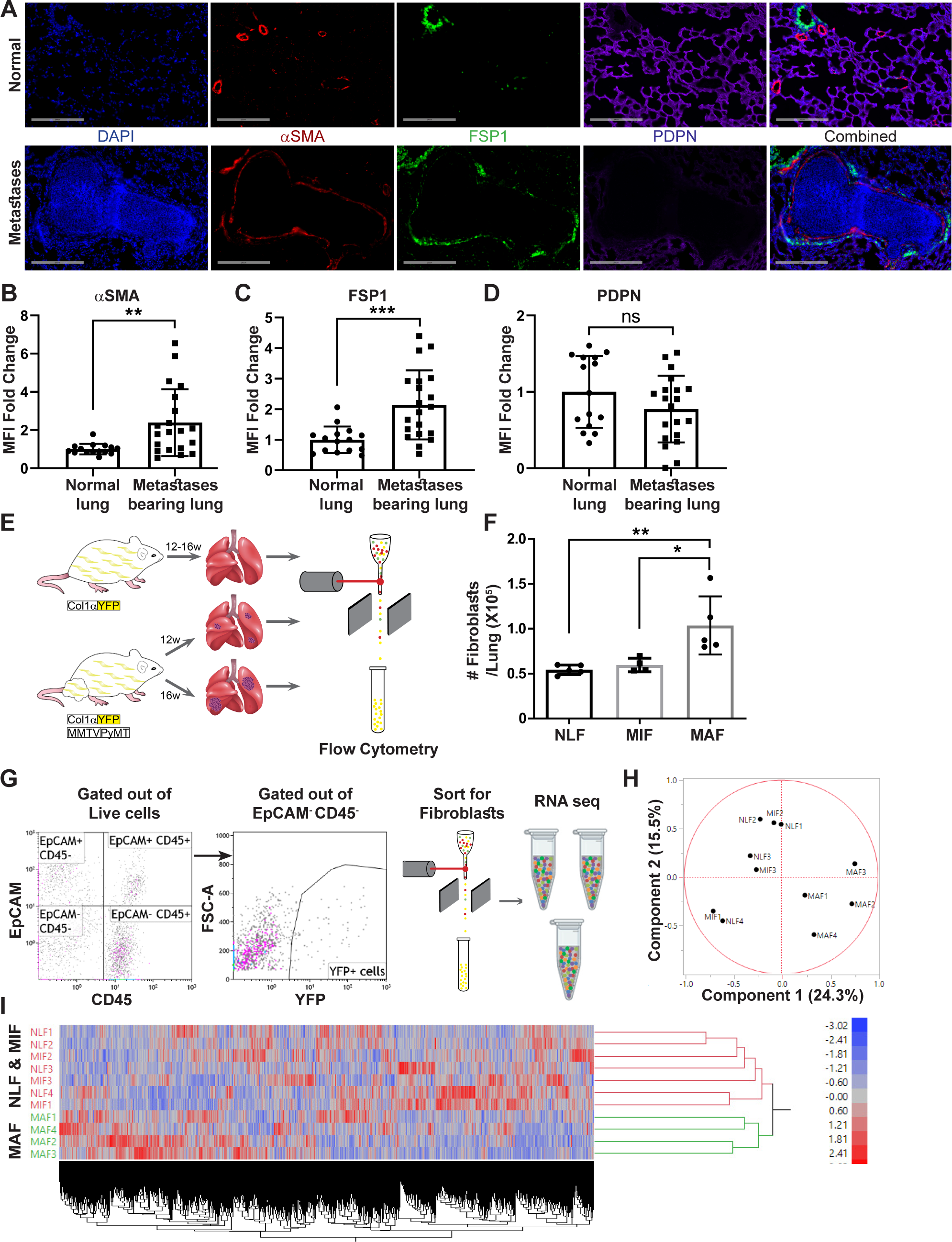
Fibroblasts are activated and transcriptionally reprogrammed in the lung metastatic niche. **(A)** Representative immunofluorescent staining of αSMA (Red), FSP1 (green) and PDPN (purple) in normal lungs from FVB/n mice (n=3), and metastases-bearing lungs from MMTV-PyMT mice (n=4). Scale bar: 200µM **(B-D)** Quantification of mean fluorescent intensity (MFI) in 5 fields of view (FOV) per mouse of staining shown in (A). **(E)** Workflow illustration of fibroblast isolation (CD45^-^EpCAM^-^YFP^+^) from normal FVB/n;*col1a1*-YFP mice (NLF), and of micro-or macrometastases-associated fibroblasts from MMTV-PyMT;*Col1a1*-YFP mice (MIF and MAF). **(F)** Quantification of number of fibroblasts per lung, based on flow cytometry analysis *p<0.05, **p<0.01. Data are represented as mean ± SD, n=5. **(G)** Flow cytometry gating strategy for isolation of fibroblasts prior to RNA-sequencing. **(H-I)** Principal Component Analysis (PCA) (H) and hierarchical clustering (I) of 11,115 protein coding genes identified in RNA-seq.

We therefore set out to characterize the changes in fibroblasts at the metastatic niche during the formation of spontaneous lung metastases. To enable visualization, tracking, and isolation of fibroblasts, we established a transgenic mouse model of breast cancer with fibroblast-specific reporter genes: transgenic mice that express the fluorescent reporter YFP under the Collagen-1α promoter (*Col1a1-*YFP) were crossed with MMTV-PyMT mice to create PyMT;*Col1a1-*YFP transgenic mice, in which all fibroblasts are fluorescently labeled ^26^. Flow cytometric analysis of normal lungs as compared with lungs of tumor-bearing mice revealed significantly increased numbers of fibroblasts in macro-metastatic lungs (Figure 1E,F). Thus, fibroblasts are both activated and increase in numbers in the metastatic microenvironment of breast cancer lung metastasis.

To analyze the transcriptional reprograming of activated fibroblasts at the lung metastatic niche we performed unbiased profiling by RNA-seq of fibroblasts isolated from lungs of PyMT;*Col1a1-*YFP transgenic mice at distinct metastatic stages, compared with fibroblasts isolated from normal lungs of *Col1a1-*YFP mice. To explore the temporal changes in functional gene networks, we profiled fibroblasts (EpCAM^-^CD45^-^YFP^+^ cells) isolated from normal lungs, and from lungs with micro- or macrometastases (Figure 1G). Micrometastases were defined by the presence of tumor cells in lungs, where no lesions were detectible macroscopically or by CT imaging.

Initial data analysis indicated that fibroblasts isolated from lungs with macrometastases (macrometastases-associated fibroblasts-MAF) were strikingly different from NLF as well as from fibroblasts isolated from lungs with micrometastases (micrometastases-associated fibroblasts-MIF) (Figure 1H,I Supplementary Figure 1A). Notably, since fibroblasts were isolated from entire lungs, rather than from specific metastatic lesions, the MIF fraction contained a mixture of normal, non-metastases-associated fibroblasts as well as metastases-associated fibroblasts. As a result, initial data analysis did not reveal significant differences between NLF and MIF. Thus, metastases-associated fibroblasts are not only functionally activated but also transcriptionally reprogrammed.

### Transcriptome profiling of metastases-associated fibroblasts reveals dynamic stage-specific changes in gene expression

In light of these initial results we next analyzed the genes that are differentially expressed between MAF and NLF. We selected upregulated and downregulated genes based on fold change of |2|. Expectedly, hierarchical clustering based on these genes revealed that the MAF group clustered separately from NLF and MIF (Figure 2A). To better characterize the trajectory of changes in fibroblasts during metastases formation, we next compared the expression of genes that were differentially expressed between MAF and NLF to their expression in the MIF population. Interestingly, we found that the expression pattern in MIF was distinct from both the MAF and the NLF gene expression, including genes that had opposite changes in MAF vs. MIF, suggesting that they are activating a distinct transcriptional program (Figure 2B).

**Figure 2.**
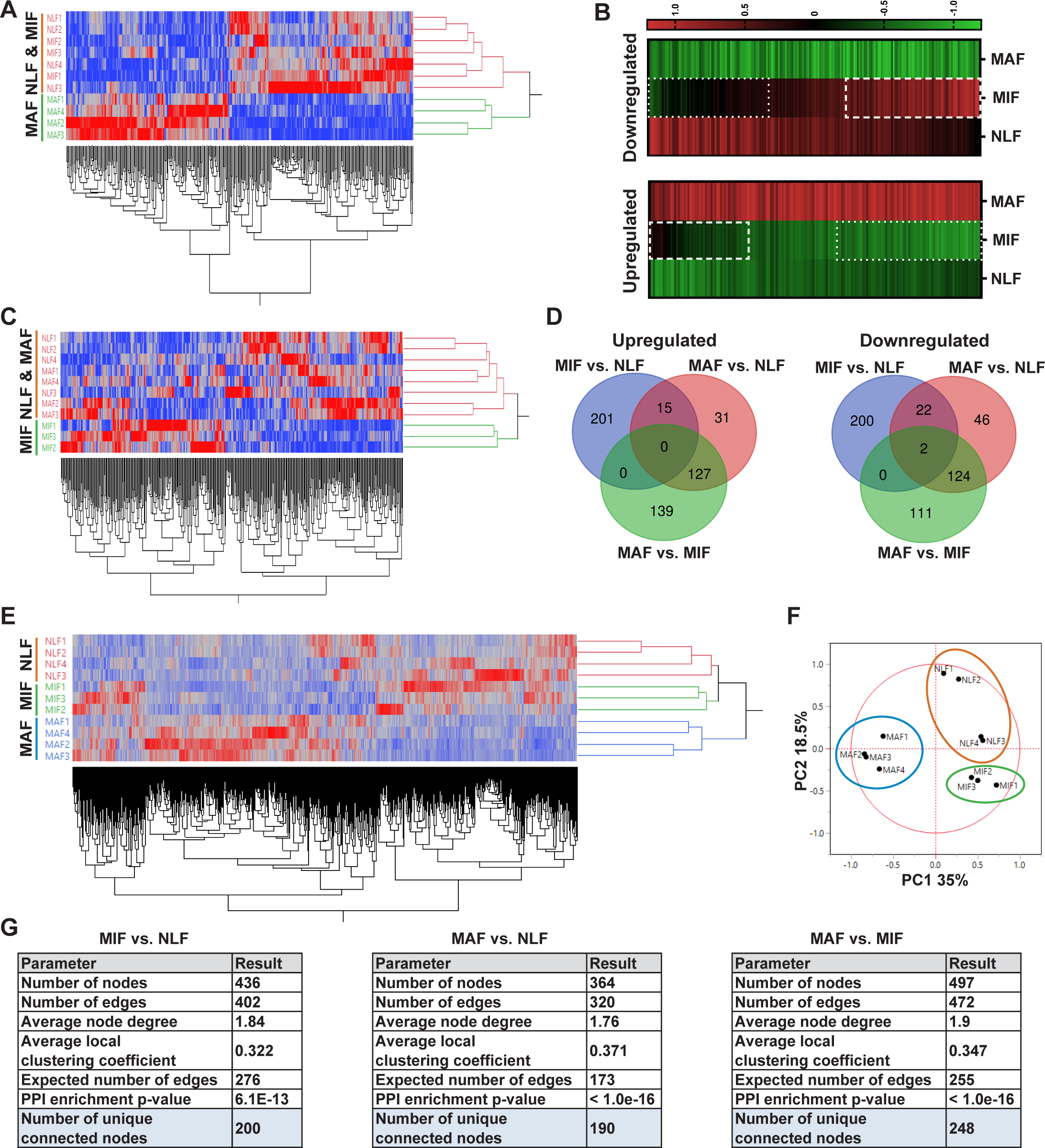
Transcriptome profiling of metastases-associated fibroblasts reveals dynamic stage-specific changes in gene expression. **(A)** Hierarchical clustering of genes upregulated or downregulated in MAF vs. NLF based on FC>|2|. **(B)** Presentation of the average Z-scored gene expression of genes differentially expression in MAF vs. NLF in all three groups: NLF, MIF and MAF. Dashed lines demarcate genes upregulated in MIF vs. NLF. Dotted lines demarcate genes downregulated in MIF vs. NLF. **(C)** Hierarchical clustering of genes upregulated or downregulated in MIF vs. NLF based on FC>|1.5|. **(D)** Venn diagram of upregulated or downregulated genes in the different comparisons. **(E**,**F)** Hierarchical clustering (E) and PCA (F) of genes upregulated or downregulated in the different comparisons (MIF vs. NLF, MAF vs, NLF, MAF vs. MIF). **(G)** Protein-protein interaction analysis of the differentially expressed genes per comparison performed in STRING platform. Interconnected genes were selected for subsequent analysis.

We therefore analyzed the differentially expressed genes in the MIF fraction separately. Since the detectible changes in micrometastases were more subtle than the changes detected in the macrometastases group, we selected these genes based on a fold change of |1.5|, to better differentiate the MIF group from NLF. Indeed, hierarchical clustering based on these differentially expressed genes confirmed that the MIF group clustered separately from both NLF and MAF (Figure 2C). Next, we selected a group of genes based on their differential expression between the MAF and MIF groups (FC≥|2|). The combination of these yielded a total of 897 genes that were differentially expressed in MIF vs. NLF, MAF vs. NLF or MAF vs. MIF. Interestingly, only a small number of these genes were shared across the different stages, suggesting again that each stage is defined by its own specific gene signature (Figure 2D). Accordingly, Principal Component Analysis (PCA) and hierarchical clustering applied on the selected gene signature dataset separated each of the metastatic stages (Figure 2E,F).

Thus, although the transcriptional changes in fibroblasts isolated from micrometastases may have been masked by the presence of normal fibroblasts in this fraction, further analyses suggested that MIF, as well as MAF activate a unique stage-specific transcriptional program.

Aiming to delineate the stage-specific gene signatures and the molecular mechanisms operative in metastases-associated fibroblasts, and to identify the most relevant functional pathways, we performed protein-protein interaction (PPI) analysis using the STRING platform ^32^ for each comparison separately. We found that per comparison, the differentially expressed genes had significantly more interactions than expected (Figure 2G, Supplementary Figure 1B-D), suggesting that they are functionally related. We therefore decided to focus our subsequent analyses on the functional roles of the subsets of differentially expressed genes that were found to be inner-connected.

### Fibroblast metastases-promoting tasks are driven by functional gene signatures related to stress response, inflammation, and ECM remodeling

We next asked whether the changes in the different metastasis-associated fibroblast subpopulations represent specific metastases-promoting tasks. To address this question, we performed further analysis of the selected genes in each stage by using the over-representation enrichment analysis of the Consensus Path DB (CPDB) platform ^33^. Our focus in these analyses was based on three different databases: GO ^34,35^, KEGG ^36,37^, and Reactome ^38^. For our analysis, we selected terms that represent biological processes enriched in at least two databases, with a relative overlap of at least 0.2 and at least 2 shared entities (Figure 3A). Data analysis revealed significant and stage-specific changes in functional pathways including cellular stress response, extracellular matrix (ECM) remodeling and inflammation (Figure 3B, Supplementary Table 1).

**Figure 3.**
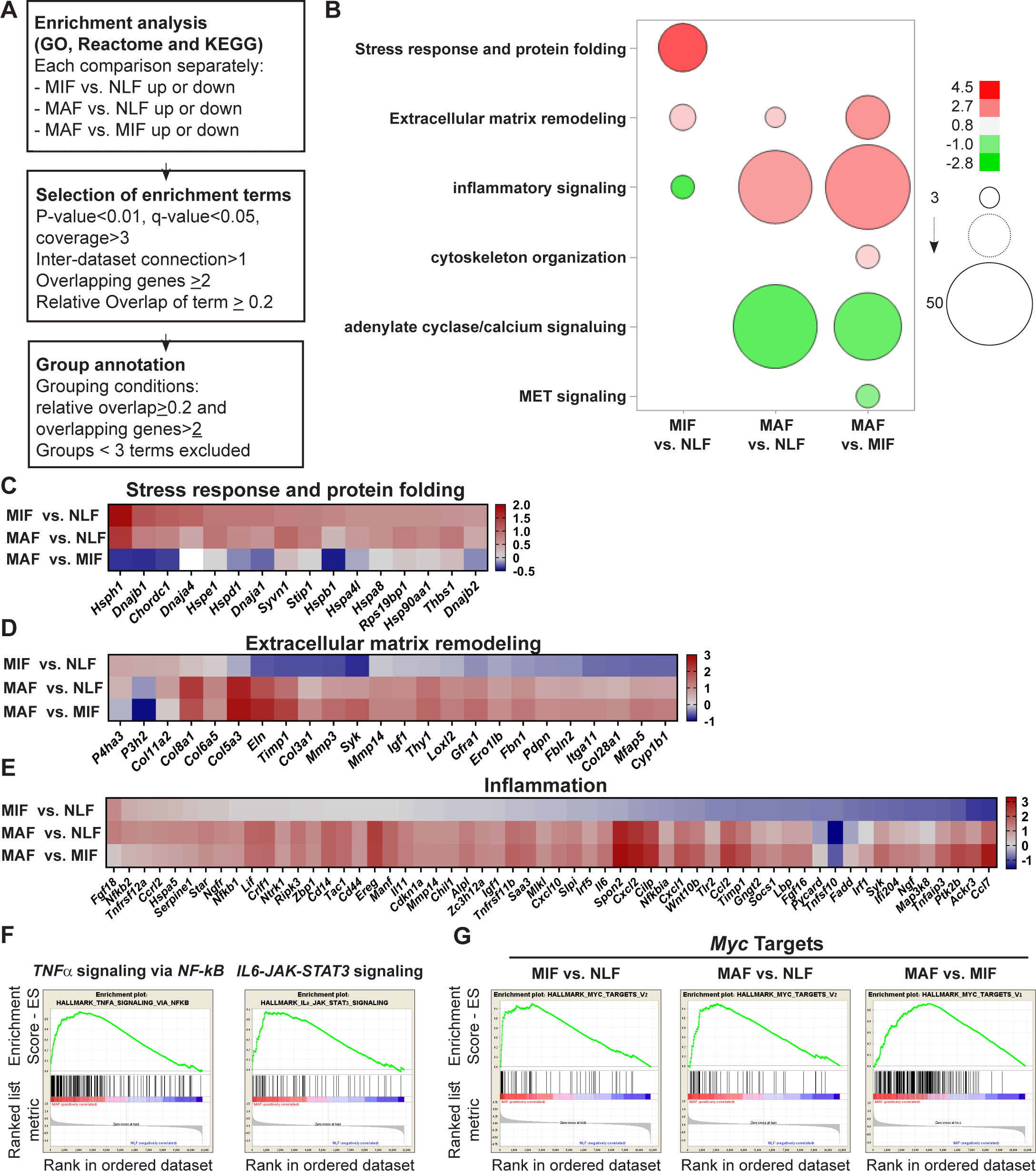
Fibroblast metastases-promoting tasks are driven by functional gene signatures related to stress response, inflammation, and ECM remodeling. **(A)** Flow chart of the pathway enrichment over-representation analyses based on GO, Reactome and KEGG using the CPDB platform. **(B)** Bubble graph heatmap based on the number of specific enrichment terms and their average Log-transformed q-value per group. Circle sizes denote number of terms included in a group, color indicates the average Log-transformed q-value. Enrichments based on downregulated genes are presented as negative values. **(C-E)** Heat maps of gene expression fold-change presenting genes in selected group annotations. Fold change was log_2_ transformed for presentation. Only genes found in at least 2 different terms are presented. (C) “Stress response and protein folding” enriched genes. (D) “Extracellular Matrix remodeling” enriched genes. (E) “Inflammatory signaling” and/or “Cytokine and Chemokine activity” enriched genes. **(F)** Gene Set Enrichment Analysis (GSEA) for hallmark datasets upregulated in MAF vs. NLF related to inflammatory signaling, FDR<0.05, NES>2. **(G)** GSEA results for “*Myc* targets” hallmark dataset that were upregulated in all comparisons. FDR<0.05 NES>2.

Interestingly, we found that gene expression signatures in fibroblasts isolated from the micro-metastatic stage were highly and specifically enriched for functions related to cellular response to stress, including *HSF1* activation, heat shock response and response to unfolded protein (Supplementary Table 1). Upregulated genes in MIF that were related to stress and protein folding included several heat shock proteins: *Hspa8, Hsp90aa1, Hspd1, Hspe1* and others (Figure 3C). Of note, while in the pathway analysis the stress response pathway was not significantly enriched in MAF (Figure 3B), detailed fold change analysis indicated that while this pathway is most upregulated in MIF, it remains upregulated in MAF compared to normal fibroblasts (Figure 3C). ECM remodeling terms were enriched in both MIF and MAF (Figure 3B), indicating the central importance of ECM modifications in facilitating metastasis. Notably, while ECM remodeling was operative throughout the metastatic process, the specific genes related to ECM remodeling in the different metastatic stages were distinct (Figure 3D).

Gene expression signatures in fibroblasts isolated from macrometastases were highly enriched for inflammation-related pathways (Figure 3B, Supplementary Table 1). Indeed, analysis of enriched pathways revealed that genes related to inflammation including many chemokines and cytokines were upregulated specifically in MAF (Figure 3E). Taken together, these findings imply that metastases-associated fibroblasts assume distinct functional roles during the process of lung metastasis.

Encouraged by these findings, we next set out to obtain further insights on functional pathways that were modified in fibroblasts isolated from different metastatic stages. To that end, we performed Gene Set Enrichment Analysis (GSEA) ^39^. We focused our analysis on the H collection: Hallmark gene sets that summarize specific well-defined biological states or processes based on multiple datasets ^40^. Similarly to the results obtained in our previous analyses, we found that functions related to inflammatory responses, including TNFα and IL-6 signaling were enriched in MAF (Figure 3F, Supplementary Table 2). Interestingly, we found that *Myc* target genes were the most highly and significantly enriched in both metastatic stages (Figure 3G, Supplementary Table 2), suggesting that this transcription factor may play a central role in the functional molecular co-evolution of metastases-associated fibroblasts.

Taken together, these findings imply that the transcriptome of lung fibroblasts is rewired during metastatic progression, driving changes in the expression of distinct molecular pathways. Moreover, the transcriptional changes in ECM remodeling and stress response, which are potential metastases-promoting tasks, are evident at early stages of metastases formation, suggesting that fibroblasts play an important role already at the onset of the metastatic process.

### Multiple gene network analyses identify Myc as a central transcription factor in the rewiring of metastases-associated fibroblasts

To further characterize the regulatory nodes that govern the functional changes in fibroblasts, we hypothesized that these changes may be driven by transcription factors related to the pathways that were identified by the pathway and GSEA analyses (Figure 3). Analysis of transcription factors terms within the results identified five candidate transcription factors (TFs) that were enriched in at least one analysis and in at least one metastatic stage: *Hif1a, Hsf1, Myc, Nfkb1 and Stat3* (Supplementary Table 3).

We next examined in how many different comparisons each TF was enriched. We found that *Hsf1* was only enriched in the micro-metastatic stage vs. normal lungs, and *Hif1a* was enriched only in the macro-metastatic stage vs. normal lungs. *Nfkb1* and *Stat3* were enriched in the macro-metastatic stage, compared with both normal and micro-metastases. Notably, only *Myc* was enriched in all three comparisons (Supplementary Table 3).

To rank these TFs, we performed knowledge-based multiple analyses examining their centrality in the selected gene signatures in each comparison (Supplementary Table 4). We examined the protein-protein interactions (PPIs) of these TFs utilizing the STRING platform, and counted the number of direct connections of each TF with the metastases-associated gene signatures. In MAF gene signature, *Stat3* had the largest number of connections, closely followed by *Myc*. In MIF gene signature, *Myc* had the largest number of connections (Figure 4A, orange). In addition to STRING, we examined PPIs using ANAT (Advanced Network Analysis Tool) ^41^. In this platform, the inference is based on setting all the candidate TFs as anchors and the selected genes as targets in a network of PPI, and searching for a putative compact sub-network that connects them. We analyzed the results according to three parameters: the number of direct connections of each TF, the characteristic path length to all nodes (including non-directly related), and network centralization. Analysis of the results revealed that *Myc* had the largest number of direct connections in all comparisons, and is overall connected to the fibroblast metastases-associated gene signatures with the shortest path and with the highest centrality in all comparisons (Figure 4A, yellow, Figure 4B, Supplementary Figure 2). These results suggested that *Myc* plays a central role in mediating the transcriptional rewiring of fibroblasts in the lung metastatic niche across the different stages.

**Figure 4.**
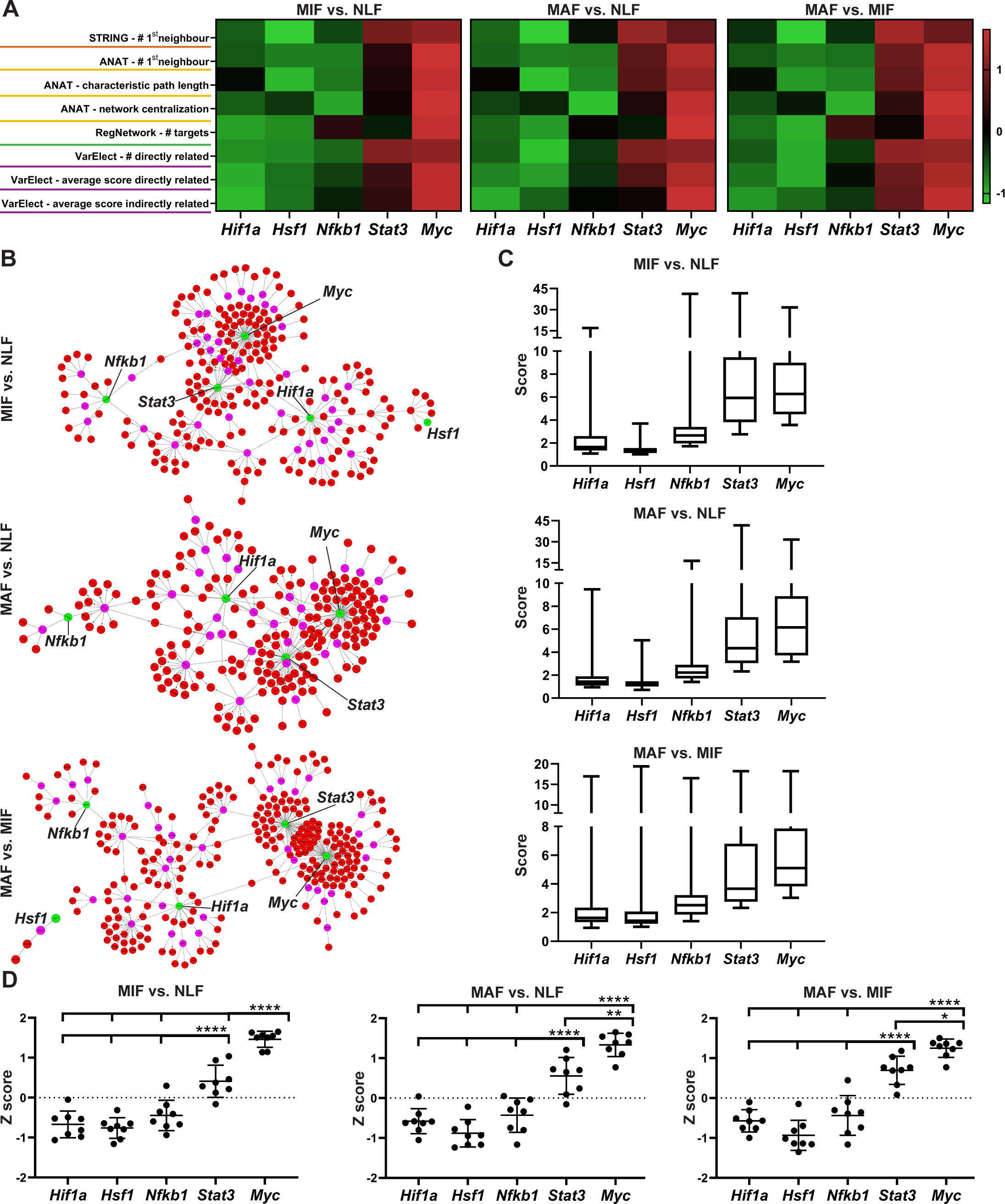
Multiple gene network analyses identify Myc as a central transcription factor in the rewiring of metastases-associated fibroblasts. **(A)** Heat maps of ranking parameters and analyses performed per each comparison to identify the centrality of five candidate transcription factors (TF): *Hif1a, Hsf1, Myc, Nfkb1, Stat3*,. Orange – STRING PPI analysis results. Yellow - ANAT pathway analysis results. Green - RegNetwork analysis of connectivity between target genes and TFs. Purple - VarElect analysis results. **(B)** Representative ANAT protein-protein network, using all TFs as anchors (green) and the stage-specific signature as target genes (red). Only interaction confidence >0.6 are presented. **(C)** Box Plot of VarElect scores for directly related genes to each TF (Presenting top 50 per TF). **(D)** Z-score Graphs of the results described in (A), *p<0.05, **p<0.01, *** p<0.001, ****p<0.0001, one-way ANOVA with Tukey correction for multiple comparisons. Data are presented as mean ± SD.

We next examined the specific connection of each TF as a regulator in the metastases-associated gene network. To that end, we utilized the RegNetwork tool ^42^, a knowledge-based database of gene regulatory networks. We found that *Myc* had the greatest number of targets in all comparisons, followed by *Stat3* and *Nfkb1* (Figure 4A, green). Finally, we analyzed the correlation of the metastases-associated gene network with each candidate TF using the VarElect tool ^43^. This tool enables prioritization of genes related to a specific query term by using a direct and indirect relatedness score. We analyzed the scores of the stage-specific signature genes with each candidate TF, and the number of directly related genes. The TFs were ranked based on the number and average score for the directly related genes, and the average score of the indirectly related genes. In agreement with previous analyses, *Myc* had the highest number of connections and the highest average score for both directly and indirectly related genes in all comparisons (Figure 4A, pink, Figure 4C). To consolidate these comprehensive gene network analyses, we performed a comparative analysis on the TF bioinformatics measurements listed in Figure 4a. The results indicated that *Myc* achieved significantly higher scores than all other TFs in all three gene signatures (Figure 4D). These results implicate the centrality of *Myc* in the dynamic transcriptional changes that govern the function of metastases-associated fibroblasts in lung metastasis.

### Myc is a central regulator in metastases-associated fibroblasts and contributes to their acquisition of tumor-promoting traits

*Myc* (myelocytomatosis oncogene) is a transcription factor involved in many biological processes, including cell growth and proliferation, cell stemness, and metabolism. *Myc* is deregulated in many human cancers, and is known to play an important role in the pathogenesis of cancer, particularly in cancer cells ^44,45^.

To validate the ranking results, we analyzed by qRT-PCR the expression of *Myc* in fibroblasts isolated from normal lungs, or from lungs with micro- and macrometastases. Analysis of the results indicated that *Myc* is significantly upregulated in macrometastases-associated fibroblasts (Figure 5A). In addition, we assessed the expression of central *Myc* targets that we found to be upregulated in metastases-associated fibroblasts, including *Hspe1, Hsp90aa1, Odc1* and *Fosl1* ^46,47^. The results indicated that these *Myc* targets were upregulated in fibroblasts isolated from lungs with metastases (Figure 5B). qRT-PCR results validated our findings in the RNA-seq data, in which *Hsp90aa1* was upregulated in MIF, whereas the other *Myc* targets were upregulated in MAF (Figure 5B). To elucidate the functional importance of *Myc* in mediating lung fibroblast reprogramming, we targeted its expression by a specific *Myc* targeting siRNA in primary lung fibroblasts. Abrogation of *Myc* expression by siMyc resulted in significant inhibition of *Myc* expression as compared with control fibroblasts (Figure 5C). Importantly, control fibroblasts highly upregulated the expression of *Myc* in response to tumor cell secreted factors (Fig, 5C, left bars), while *Myc* inhibition abrogated the upregulation of *Myc* in response to tumor cell secreted factors in activated fibroblasts (Figure 5C, right bars). We next assessed whether inhibition of *Myc* affected the expression of selected *Myc* target genes in activated lung fibroblasts. Analysis of the results indicated that targeting the expression of *Myc* significantly inhibited the expression of its target genes in response to tumor cell conditioned media (CM), indicating that the expression of *Myc* in fibroblasts is central to the upregulation of its known targets (Figure 5D). Finally, we examined the importance of *Myc* for functional reprogramming of fibroblasts. Fibroblasts at the primary tumor site were previously shown to be reprogrammed by tumor cell-derived paracrine signaling ^25,48^. We therefore first asked whether fibroblasts at the metastatic microenvironment are similarly activated in response to tumor-secreted factors. Incubation of isolated primary lung fibroblasts with CM from Met-1, a PyMT-derived breast carcinoma cell line ^49^, or from 4T1 cells, a model of triple-negative breast cancer, indicated that tumor-derived factors activated multiple CAF-associated tasks including enhanced motility in wound healing assay (Supplementary Figure 3A-D) and increased contraction of collagen gel matrices (Supplementary Fig 3E-G). Thus, normal lung fibroblasts are reprogrammed by signaling from breast cancer cells, resulting in acquisition of tumor-promoting properties. To test whether activation of Myc in lung fibroblasts contributes to their acquisition of CAF characteristics, we performed wound healing assays and collagen contraction assays with tumor-activated lung fibroblasts that were transfected with siMyc or with siCtrl. We found that siMyc fibroblasts were less contractile and exhibited significantly attenuated migration capacity as compared with controls (Figure 5E-H). Taken together, our findings imply that *Myc* has a central role in enhancing fibroblast activation and in mediating their acquisition of metastasis-promoting functions.

**Figure 5.**
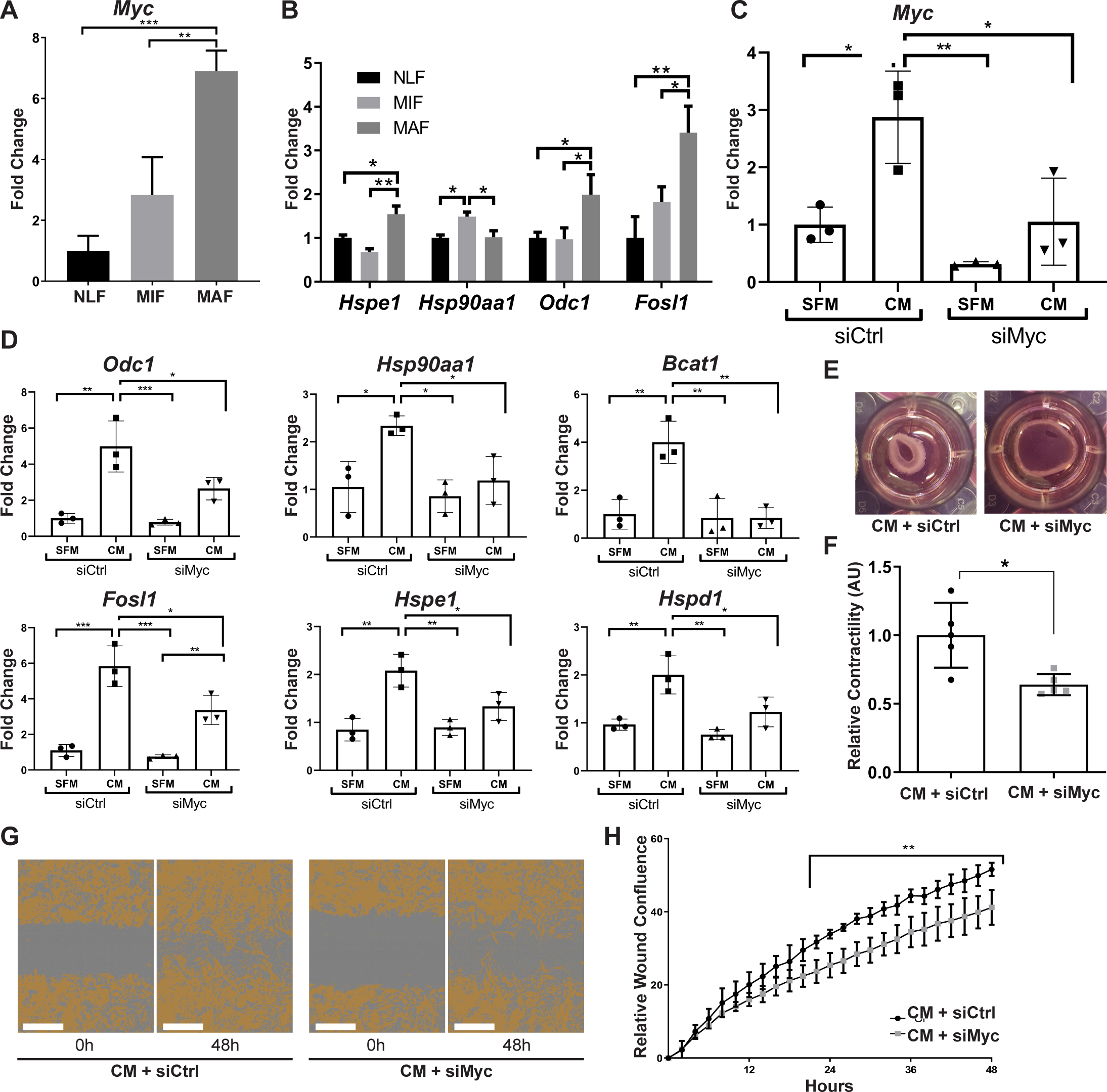
Myc is a central regulator in metastases-associated fibroblasts and contributes to their acquisition of tumor-promoting traits. **(A)** qRT-PCR analysis of *Myc* expression in sorted NLF, MIF and MAF. **p<0.01, Data are represented as mean ± SD, n=3 per group. **(B)** qRT-PCR analysis in sorted NLF, MIF and MAF. Relative expression of *Myc* target genes found to be differentially expressed in RNA-seq. *p<0.05, Data are presented as mean ± SD, n=3 per group. **(C)** *Myc* targeting by siRNA: *Myc* expression in NLF transfected with siRNA targeting *Myc* or with control siRNA (siMyc or siCtrl). Following transfection, cells were incubated with SFM or with Met-1 CM supplemented with the same siRNA for additional 24h. Data are presented as mean ± SD, n=3. **(D)** qRT-PCR analysis of *Myc* targets following treatment as in (C). Data are represented as mean ± SD, n=3. **(E**,**F)** Representative images and quantification of collagen contraction assay of fibroblasts transfected with siMyc or siCtrl, incubated with Met-1 CM. *p<0.05, data are represented as mean ± SD, n=5. **(G**,**H)** Representative images and quantification of scratch closure assay of NLF transfected with siMyc or siCtrl and incubated with Met-1 CM. Scale bar: 400µm. Two-way ANOVA with multiple comparisons, ***p<0.001, data are presented as mean ± SD, n=5.

### High expression of MYC and its downstream target genes is associated with worse survival in human breast cancer

Encouraged by these findings, we next asked whether stromal activation of *MYC* and its downstream targets is operative in human breast cancer. There are currently no available transcriptomic datasets of lung metastases, and we therefore analyzed patient data from breast tumors utilizing a publicly available dataset ^50^. Since we showed that *MYC* is a central regulator of fibroblast rewiring during metastatic progression in mice, we asked whether *MYC* is similarly upregulated in the stromal compartment of human breast cancer. Importantly, analysis revealed that *MYC* is upregulated in breast cancer stroma in correlation with disease progression, as reflected by pathological grade: expression of *MYC* was elevated in the stroma of grade 3 tumors, compared with stroma isolated from more differentiated tumors (Figure 6A). Interestingly, *NFKB1* and *STAT3* did not exhibit this grade-dependent trend of expression (Figure 6B,C). To further assess whether the upregulation of stromal *MYC* and its target genes is operative in the stromal compartment of human breast tumors, we compared the expression of *MYC* with the expression of its target genes in human breast cancer patients. Target genes were selected based on their upregulation in metastases-associated fibroblasts. We found that stromal expression of *MYC* was positively correlated with stromal expression of multiple target genes (Figure 6D). Notably, among the *MYC* downstream target genes that were positively correlated with its expression in human patients, were several of the genes that were also validated in murine lung fibroblasts: *HSP90AA1, HSPD1, ODC1* and *HSPE1* (Figure 6D, Supplementary Figure 4). Motivated by these findings we assessed whether expression of *MYC* targets is related to survival in human breast cancer. We stratified patients based on their expression of *MYC* targets that were upregulated in micrometastases or macrometastases and analyzed the correlation between their level of expression and patient survival. Analysis revealed that higher expression of MYC target genes was significantly correlated with worse survival (Figure 6E). Of note, these survival data are derived from a dataset of total tumor tissue, and therefore the results are not exclusively due to stromal expression. Nevertheless, these findings indicate that upregulation of *MYC*-driven gene expression signatures that we found to be operative in metastases-associated stroma may play a role in progression and survival of human breast cancer patients. Finally, to validate our findings in human metastasis, we analyzed the expression of MYC in a cohort of breast cancer patients with lung metastasis. We found that MYC was expressed in a subset of lung metastases-associated fibroblasts (Figure 6F), suggesting that stromal upregulation of MYC plays a functional role in human lung metastasis.

**Figure 6.**
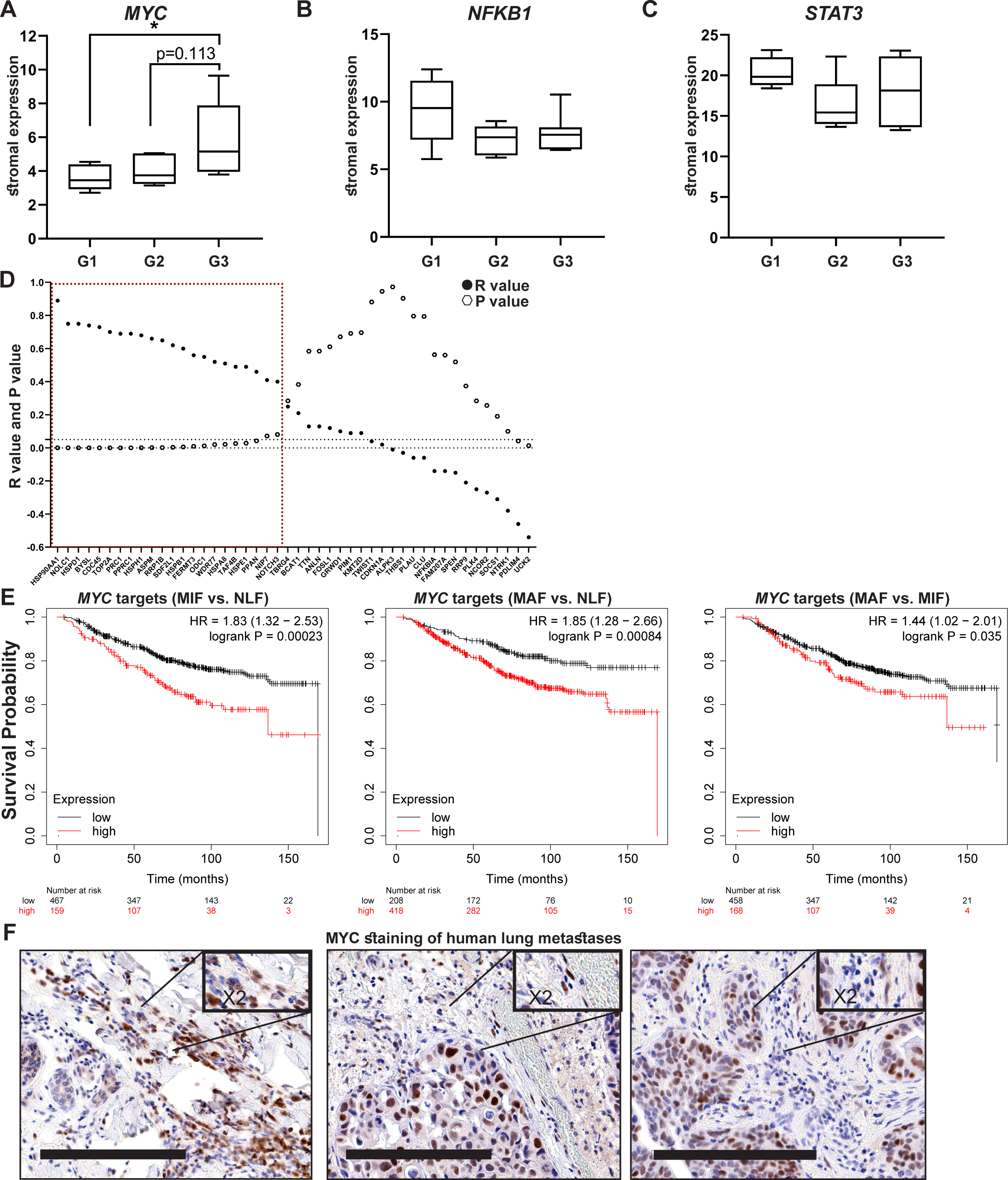
High expression of MYC and the metastases-associated fibroblast gene signature is associated with worse survival in human breast cancer. **(A-C)** Box-plots of *MYC* (A) *NFKB1* (B) and *STAT3* (C) expression in tumor associated-stroma from the GSE14548 dataset by disease grade (grade 1: G1; grade 2: G2; grade 3: G3). Data are presented as median and upper and lower quartiles ± SD. One-way ANOVA with Tukey correction for multiple comparisons, *p <0.05. **(D)** Correlations between the expression of *MYC* and selected downstream targets in tumor-associated stroma based on GSE14548. Positive correlations are marked in dotted red square. *p-value<0.05. **(E)** Survival analysis (KM plotter) based on the combined expression of Myc downstream genes upregulated at different metastatic stages. **(F)** Representative IHC staining of MYC in lung metastases of breast cancer patients (n=9). Scale bars: 200μm

Taken together, these results suggest that the activation of MYC transcriptional networks in the stroma of breast tumors plays a role in tumor progression and survival in human breast cancer.

## Discussion

In this study we set out to elucidate the dynamic changes in the stromal compartment that facilitate the formation of a hospitable metastatic niche during breast cancer metastasis to lungs. We utilized a unique model of transgenic mice that enabled unbiased isolation of fibroblasts from spontaneous lung metastasis and performed comprehensive analysis of the transcriptome of fibroblasts isolated from normal lungs, and lungs with micro- or macrometastases. By employing multiple platforms of data analysis, we integrated ontology analyses with data on protein interactions and functional pathways from knowledge-based databases to identify the relevant and stage-specific gene signatures that imply functional tasks of metastases-associated fibroblasts.

Importantly, we performed the analysis on fibroblasts isolated directly from fresh tissues, with no additional culture step that may affect gene expression. Our findings indicated that ECM remodeling programs were instigated early in micrometastases, and persisted to be functional throughout metastatic progression, while other signaling pathways were activated in a stage-specific manner. Activation of the cellular stress response was associated with micrometastases, and inflammatory signaling was instigated in fibroblasts isolated from advanced metastases, suggesting that fibroblasts are transcriptionally dynamic and plastic, and that they adapt their function to the evolving microenvironment (Figure 7).

**Figure 7.**
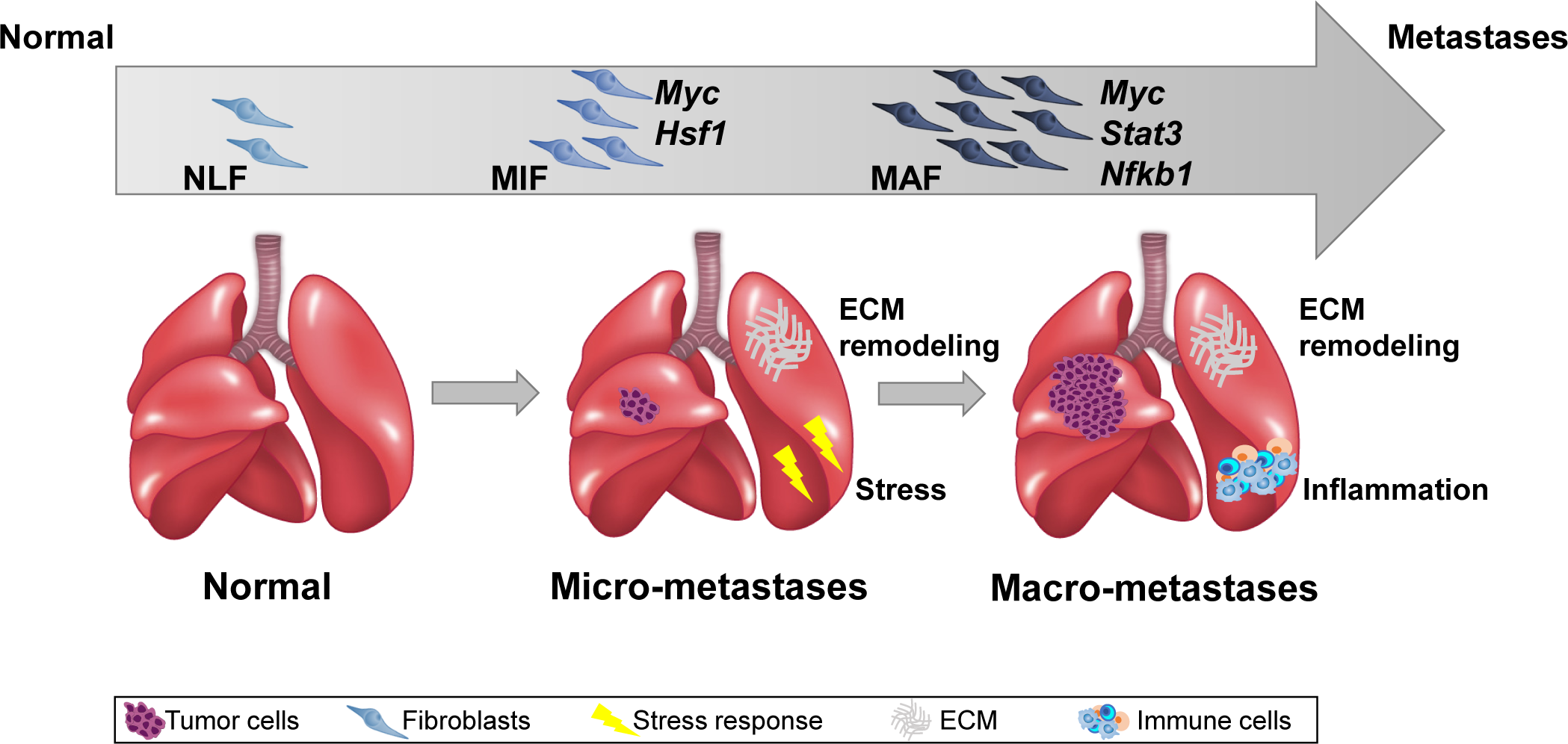
Summary scheme: The co-evolution of lung fibroblasts at the metastatic microenvironment is driven by stage-specific transcriptional plasticity that activates growth-promoting tasks including stress response, ECM remodeling and instigation of inflammatory signaling.

Initial analysis of the RNA-seq data revealed distinct gene signatures associated with advanced metastatic disease. By performing step by step analysis, a unique gene signature was revealed for early metastatic disease as well. Moreover, utilizing a combination of analyses platforms, we unraveled multiple pathways operative in fibroblasts in different metastatic stages, relying not only on altered gene expression but also on functional role and interaction of genes.

Interestingly, this multi-layered analysis indicated that fibroblasts isolated from micrometastases instigated the expression of genes related to cellular response to stress, including the transcriptional regulator *Hsf1. Hsf1* was previously shown to be upregulated in CAFs in breast and lung cancers and to drive a stromal tumor-promoting transcriptional program that correlated with worse prognosis ^51^. Moreover, *Hsf1* was recently implicated in activating stromal expression of Dickkopf-3 (*Dkk3*) in mammary fibroblasts, which is a central modulator of their pro-tumorigenic activity, thus linking *Hsf1* with reprogramming of stromal fibroblasts to CAFs ^52^. Our findings expand these observations to the metastatic microenvironment, and show that activation of *Hsf1* transcriptional regulation in fibroblasts occurs during the early stages of metastasis and thus may play a role in instigating tumor-promoting tasks in metastases-associated fibroblasts.

In addition to stress response, our findings indicated that ECM remodeling is a central task of metastases-associated fibroblasts throughout the metastatic cascade. Indeed, ECM components and remodeling were demonstrated to facilitate breast cancer metastasis to lungs, and pancreatic cancer metastasis to liver ^13,14,16,53-55^. We show that transcriptional rewiring of fibroblasts to mediate tasks such as collagen synthesis and ECM organization is a central function of metastases-associated fibroblasts, which is instigated early during the metastatic process and persists during advanced metastatic disease.

Notably, analyzing the central pathways in fibroblasts that were isolated from advanced metastases, indicated that metastases-associated fibroblasts upregulated pro-inflammatory pathways including multiple cytokines and chemokines. CAFs are known to play a central role in mediating tumor-promoting inflammation at the primary tumor site ^56^. Importantly, in recent years activation of inflammation has also emerged in shaping of the metastatic microenvironment ^10,11,18^, but the role of fibroblasts in mediating inflammation at the metastatic site was not previously shown. Our findings implicate CAF-derived inflammatory signaling in promoting the growth of lung metastases. Further studies are required to determine which pro-inflammatory factors upregulated in metastases-associated fibroblasts are functionally important in formation of a hospitable metastatic niche in lungs.

We further characterized the molecular mechanisms operative in metastases-associated fibroblasts, by identifying the central transcription factors that drive the metastases-associated gene programs upregulated in lung fibroblasts. Our analyses revealed several central regulators that are operative in metastases-associated fibroblasts, including the well-known modulators of CAF activity *Nfkb1* ^23,24^ and *Stat3* ^57,58^.

Surprisingly, the most prominent regulator in the metastasis-associated fibroblasts network was the transcription factor *Myc*. While the importance of *Myc* in promoting cell transformation and tumorigenesis is well established ^45^, its role in the tumor stroma is largely uncharacterized. *Myc* expression in tumor cells was shown to drive an inflammatory and immunosuppressive microenvironment ^59^. Moreover, the expression of *Myc* in the stromal compartment was suggested to mediate metabolic and transcriptional reprogramming of fibroblasts ^60,61^. Our study identifies *Myc* as a central regulator in the transcriptional plasticity of metastases-associated fibroblasts. Indeed, inhibition of *Myc* attenuated tumor promoting functions of fibroblasts, confirming that *Myc* functionally contributes to their acquisition of tumor promoting traits. Importantly, validation of these findings in human breast cancer patients revealed that stromal expression of *Myc* and its downstream genes is highly correlated with worse survival in breast cancer patients. Stromal gene expression was previously found to be associated with bad prognosis in colon cancer ^62^. Our findings implicate activation of *Myc* and stromal gene expression in breast cancer patient survival. Taken together, these findings indicate that in addition to its known role in driving carcinogenesis in tumor cells, *Myc* functions in stromal rewiring in the tumor microenvironment in both primary tumors and metastases of breast cancer.

In summary, we show that integration of multiple analytical platforms of gene expression, connectivity and function provided a powerful insight on functional and temporal regulation of the dynamic transcriptome of fibroblasts in lung metastasis. We uncovered central molecular pathways that drive the activation of growth-promoting tasks in fibroblasts via known regulators of CAF tumor-promoting activities including *Myc*, a novel regulator of fibroblast metastases-promoting tasks. Our findings elucidate for the first time the dynamic transcriptional co-evolution of fibroblasts during the multi-stage process of breast cancer metastasis.

## Methods

### Mice

All experiments were performed using 6-8 weeks old female mice, unless otherwise stated. All experiments involving animals were approved by the Tel Aviv University Institutional Animal Care and Use Committee. FVB/n *Col1a1*-YFP mice were a kind gift from Dr. Gustavo Leone. FVB/N-Tg (MMTV-PyMT) 634Mul/J were backcrossed with FVB/n;*Col1a1*-YFP mice to create PyMT;*Col1a1*-YFP double-transgenic mice as described previously ^26^. Non-transgenic Balb/c mice were purchased from Harlan, Israel. All animals were maintained within the Tel Aviv University Specific Pathogen Free (SPF) facility.

### Cell cultures

Cancer cell lines: Met-1 mouse mammary gland carcinoma cells were a gift from Prof. Jeffrey Pollard. Met-1 cells were plated on 100mm plastic dishes and cultured with DMEM medium supplemented with 10% FCS, 1% penicillin-streptomycin and 1% Sodium-pyruvate (Biological Industries). 4T1 mouse mammary cell lines were obtained from the laboratory of Dr. Zvi Granot. 4T1 cells were plated on 100mm plastic dishes and cultured with RPMI medium supplemented with 10% FCS, 1% penicillin-streptomycin and 1% Sodium-pyruvate (Biological Industries).

Primary lung fibroblasts cultures: Lungs were isolated from 6-8 weeks old FVB/n female mice or Balb/C female mice. Single cell suspensions were prepared as previously described ^63^. Single cell suspensions were seeded on 6-well plates pre-coated with Rat tail collagen (Corning; 354236). Cells were grown in DMEM media supplemented with 10% FCS, and maintained at 37°C with 5% CO_2_.

### Conditioned media

Tumor cell conditioned media (Met-1 CM or 4T1 CM): cells were cultured as described above. When cells reached 80% confluency, plates were washed twice with PBS and fresh serum free medium (SFM) was applied. After 48h, medium was collected, filtered through 0.45μm filters under aseptic conditions, flash-frozen in liquid nitrogen and stored at −80°C. SFM supplemented as above was used as control.

Normal lung fibroblasts (NLF) or Activated lung fibroblasts (ALFs) conditioned media: NLF were plated as described above. CM was prepared by incubating NLF with either SFM (for NLF CM) or tumor cell CM (for ALF CM) for 24 hours. After 24h, plates were washed twice with PBS and cells were incubated for additional 24h with fresh SFM. After 48h, medium was collected, filtered through 0.45μm filters under aseptic conditions, flash-frozen in liquid nitrogen and stored at −80°C.

### Scratch assay

NLF were plated in a 96-well IncuCyte® imageLock plate (Essen BioSciense). SFM was applied for 16h. Wells were then washed three times with PBS and a scratch was made using the IncuCyte® WoundMaker (EssenBiosciense). Wells were washed three times with PBS and cancer cell CM or SFM were applied. The plate was placed in the IncuCyte® system (Essen BioSciense) for 48 hours. Images were analyzed using the IncuCyte® software. Inhibition of proliferation was performed by adding 20µg/ml mitomycin C (Sigma Aldrich; M4287) to all wells during the scratch closure time.

### Collagen contraction

NLF were plated as mentioned above and incubated with SFM for 16h. Following, Cells were detached from dishes with trypsin and counted. A total of 1.5×10^5^ fibroblasts were suspended in a medium and collagen mixture [cancer cell CM or SFM mixed with High Concentration Rat Tail Collagen, type 1 (BD bioscience)], and allowed to set at 37°C for 45 min. tumor cell CM or SFM were applied, gels were released and incubated for 24 hours. Gels were photographed at various time points. ImageJ software was used to measure gel area and assess collagen contraction.

### Migration assay

Met-1 (5×10^4^) cells were placed into the upper chamber of 24 Transwell inserts, with pore sizes of 8µm, in 300µl NLF CM or ALF CM. Following 24h incubation, the upper side of the apical chamber was scraped gently with cotton swabs to remove non-migrating cells, fixed with methanol and stained with DAPI. Migrated cells were documented under a fluorescence microscope. ImageJ software was used to quantify migration.

### Immunostaining

To prepare formalin-fixed paraffin-embedded (FFPE) blocks lungs were harvested, shortly washed in PBS and incubated for 8–12 h in 10% formalin (Electron Microscopy Sciences) and transferred through ascending dilutions of ethanol before embedment in paraffin. Serial sections were obtained to ensure equal sampling of the examined specimens (5–10 μm trimming). FFPE sections from mouse lungs were deparaffinized, and incubated in 10% Neutral buffered formalin (NBF prepared by 1:25 dilution of 37% formaldehyde solution in PBS) for 20 minutes in room temperature, washed (with PBS) and then antigen retrieval was performed by microwave (2 min full power, 1000W, then 10 min at 30% of full power) with citrate buffer (pH 6.0; for αSMA and PDPN) or with Tris-EDTA buffer (pH 9.0; for S100A4). Slides were blocked with 10% BSA + 0.05% Tween20 and the antibodies listed in the table below were diluted in 2% BSA in 0.05% PBST and used in a multiplexed manner with the OPAL reagents, each one O.N. at 4° C. The OPAL is a stepwise workflow that involves tyramide signal amplification. This enables simultaneous detection of multiple antigens on a single section by producing a fluorescent signal that allows multiplexed immunohistochemistry, imaging and quantitation (Opal Reagent pack and amplification diluent, Akoya Bioscience). Briefly, following over-night incubation with primary antibodies, slides were washed with 0.05% PBST, incubated with secondary antibodies conjugated to HRP for 10 min, washed again and incubated with OPAL reagents for 10 min. Slides were then washed and microwaved (as describe above), washed, stained with the next first antibody or with DAPI in the end of the cycle and mounted. We used the following staining sequences: αSMA, S100a4, PDPN. Each antibody was validated and optimized separately, and then multiplexed immunofluorescence (MxIF) was optimized to confirm that signals were not lost or changed due to the multistep protocol. The slides were scanned at X20 magnification using the Leica Aperio VERSA slide scanner. Quantitative analyses of fluorescence intensity were performed using ImageJ Software.

### Flow cytometry analysis and cell sorting

Single cell suspensions of Lungs isolated from FVB/n;*Col1α1*-YFP or PyMT;*Col1α1*-YFP mice were stained using the following antibodies: anti-EpCAM-APC (eBioscience, 17-5791), anti-CD45-PerCP-Cy5.5 (eBioscience; 45-0451), anti-CD31-PE-Cy7 (eBioscience; 25-0311). DAPI was used to exclude dead cells (Molecular Probes; D3571). Flow cytometric analysis was performed using CytoFLEX Flow Cytometer (Beckman Coulter). Cell sorting was performed using BD FACSAria II or BD FACSAria Fusion (BD bioscience). Data analysis was performed with the Kaluza Flow Analysis software (Beckman Coulter).

### RNA isolation and qRT-PCR

RNA from sorted cells was isolated using the EZ-RNAII kit (20-410-100, biological industries) according to the manufacturer’s protocol. RNA from *in vitro* experiments was isolated using the PureLink™ RNA Mini Kit (Invitrogen; 12183018A). cDNA synthesis was conducted using qScript cDNA Syntesis kit (Quanta; 95047-100). Quantitative real-time PCRs (qRT-PCR) were conducted using PerfeCTa SYBR Green Fastmix ROX (Quanta; 95073-012). In all analyses expression results were normalized to *Gusb or Gapdh* and to control cells. RQ (2^-ΔΔCt^) was calculated.

### Transfection of primary fibroblasts

NLF were cultured in DMEM supplemented with 10% FCS. At 70% confluency, cells were transfected with Accell Delivery Media (GE Dharmacon; B-005000) supplemented with 1µM Accell SMARTpool mouse *Myc* siRNA (Dharmacon; E-040813) or Accell Control Pool non-targeting siRNA (Dharmacon; D-001910) for 96h. Accell SMARTpool contains a mixture of four siRNAs targeting one gene, and provide extended duration of gene knockdown with only minimal effects on cell viability and the innate immune response. The efficiency of *Myc* siRNA knockdown was analyzed by qRT-PCR.

### RNA-seq

CD45^-^EpCAM^-^YFP^+^ Fibroblasts were isolated by cell sorting from Normal FVB/n; *Col1a1*-YFP mice (n=4), PyMT;*Col1a1*-YFP Micro-metastases bearing mice (n=3) and PyMT;*Col1a1*-YFP Macro-metastases bearing mice (n=4). Micro-metastases were defined as visible mammary tumors, the absence of visible macro-metastases and the presence of EpCAM^+^ cells in lungs. Cells were collected into Trizol LS reagent (Life Technologies; 10296-028) and RNA was isolated according to the manufacturer’s instructions. Transcriptomic sequencing of RNA was performed using NEBNext® rRNA Depletion Kit (New England Biolabs, Inc.; E6310S) and SMARTer Stranded Total RNA-Seq Kit - Pico Input (Clontech; 635005) and sequenced on the Illumina HiSeq 2500 sequencer (Illumina, Inc.) at the Technion Genome Center. Sequenced reads were aligned to the mouse genome (mm9) using TopHat2 ^64^. Gene expression counts were calculated using HTseq-count ^65^ using Gencode annotations. Only genes that got at least 20 counts in at least 3 replicate samples were included in subsequent analysis (12,105 genes). Gene expression counts were normalized using quantile normalization ^66^. Levels below 20 were then set to 20 to reduce inflation of fold-change estimates for lowly expressed genes. Preliminary differential expression analysis was carried out using DESeq2 ^67^. For subsequent analyses, only protein coding genes were included. In addition, coefficient of variance (CV) was calculated per group (NLF, MIF, MAF) and the top 1% most in-group deviated genes (top 1% CV) were excluded, leaving a total of 11,115 genes.

### Stage-specific signature analysis

The top altered genes from MAF vs. NLF were selected based on fold change (FC) ≥<u>|</u>2|. The MIF vs. NLF genes were selected based on a FC cutoff |1.5|. Data was Z-scored per gene. Venn diagram was generated using Bioinformatics & Evolutionary Genomics website (http://bioinformatics.psb.ugent.be/webtools/Venn/). All hierarchical clustering (based on Euclidian distance and average linkage) and principal component analyses were performed using JMP software version 14 and up.

#### Gene selection based on network connectivity

Each group of genes (MIF vs. NLF, MAF vs. NLF and MAF vs. MIF) were subjected to protein-protein interactions analysis using the STRING platform ^32^. The minimum confidence of interaction was defined as confidence ≥0.3 and connections based on text-mining were excluded. Groups of under 4 genes were excluded, narrowing the size of each group by ∼50%.

### Pathway enrichment

For functional annotation, pathway and enrichment analysis, each comparison was analyzed separately, to a total of 6 comparisons (MIF vs. NLF up, MIF vs. NLF down, MAF vs. NLF up, MAF vs. NLF down, MAF vs. MIF up, MAF vs. MIF down). Over-representation analysis was performed using the ConsensusPath DataBase CPDB, ^68^,^33^ (http://cpdb.molgen.mpg.de/MCPDB) platform for GO-molecular function (MF) and GO-biological process (BP), Reactome, and KEGG. Terms larger than 500 genes were excluded. Results were considered significant with a p-value<0.01, q-value<0.05 and a coverage ≥3%. To increase the specificity of the enriched terms, we compared the relative overlap and the number of shared entities between the enriched terms from the three different databases (GO, KEGG and Reactome). Selected terms with at least 2 shared entities and a relative overlap ≥ 0.2 were grouped and annotated based on a common enriched function. Groups smaller than 3 terms were excluded. These steps enabled the selection of the top ∼10% most highly and significantly connected terms.

Bubble plot heat maps were generated with averaged log transformed q-values [-Log_10_(q-value)]. For terms enriched in a group of downregulated genes, the value of the average log transformed q-value was transformed to a negative value by duplicating the average log transformed value by (−1).

Heat maps were generated per annotation group, with a [log_2_(Fold-change)] of gene expression calculated per comparison (MIF vs. NLF, MAF vs. NLF and MAF vs. MIF).

### Gene Set Enrichment Analysis (GSEA)

The GSEA Java plug-in was used to probe log-transformed normalized expression data ^39^ http://software.broadinstitute.org/gsea/index.jsp). Settings for the analysis were defined as the follows: Gene set database – Hallmark gene sets only, Number of permutations - 1000, comparisons – each separately (MIF vs. NLF, MAF vs. NLF, MAF vs. MIF), Permutation type - gene_set, minimum size - 5, maximum size - 500. Significant results were considered for False Discovery Rate (FDR) <0.05 and normalized enrichment score (NES) > |2|.

### Transcription Factor Ranking

#### Selection of transcription factors

Transcription factors (*Hif1a, Hsf1, Myc, Nfkb1, Stat3*) that were enriched in pathway enrichment and/or GSEA analyses were selected as candidates and subjected to subsequent analyses.

#### STRING

All five candidate TFs were subjected to protein-protein interaction analysis in combination with each list of stage-specific genes per comparison (upregulated and downregulated in MIF vs. NLF, MAF vs. NLF or MAF vs. MIF separately) using the STRING platform ^32^. The confidence of the interaction was defined as >0.2. For the ranking of each TF, the number of separate interactions for each TF was counted.

#### *Advanced Network Analysis Tool* (ANAT)

The ANAT application ^41^ was used as an add-in to Cytoscape (version 7 and up) software. We performed the analysis for each TF separately and for all of the TFs combined. The TFs were defined as anchors in the list, and the target genes were each list of stage-specific genes per comparison separately. An HTML report of all possible pathways between the anchor and each gene in the target genes list was generated. The minimum confidence for a connection was defined as confidence >0.2. An anchor could be connected to a target directly, or indirectly. For the ranking of each TF, we calculated several parameters of the protein-protein network: 1) The number of stage-specific genes connected with each TF directly (1^st^ neighbor); 2) The average shortest path for each TF; 3) The centrality of the network. Parameters 2 and 3 were calculated using the network analysis tool of the Cytoscape software.

#### RegNetwork

Each TF was defined as a regulator in the RegNetwork database ^42^. For ranking of each TF, the number of registered target genes from each list of stage-specific genes were counted.

#### VarElect

VarElect platform ^43^ was utilized to analyze the relation of each list of stage-specific genes per comparison separately with each TF. Each gene from the list received a score according to its relation to the TF. For the ranking of each TF, several parameters were considered: 1) the number of directly related genes; 2) The average score of related genes; 3) The average score of indirectly related genes.

#### Ranking

Ranking parameters described above were Z-scored per parameter. For “Characteristic path length” results were first transformed with a (−1) power. Statistical analysis was performed using One-Way ANOVA with Tukey correction for multiple comparisons.

### Human breast cancer data

The expression of the metastases-associated gene signature and *MYC, NFKB1* or *STAT3* were analyzed in human breast cancer stroma based on a publicly available dataset GSE14548 ^50^. Correlation analysis between *MYC* and its downstream genes derived from the metastases-associated gene signature was performed on normalized expression values using Pearson correlation. P-value below 0.05 was considered significant.

#### Kaplan-Meier analyses

Overall survival analysis was performed using KM plotter^69^ based on combined expression of *Myc* downstream genes that were upregulated in each stage. Groups cutoff was selected automatically using the KM platform. p-value below 0.05 was considered significant.

### Human MYC staining

Human patient samples were collected and processed at the Sheba Medical Center, Israel under an approved institutional review board (IRB) (3112-16). Sections stained for MYC were analyzed by an expert pathologist (Prof. Iris Barshack). Images were scanned at X20 magnification using the Leica Aperio VERSA slide scanner. Analysis of the staining was performed using ImageScope software.

### Statistical analysis

Statistical analyses were performed using GraphPad Prism software and JMP pro 14 and 15 software. For two groups, statistical significance was calculated using t-test with Welch correction. For more than two comparisons, One-Way ANOVA with Tukey correction for multiple comparisons was applied. All tests were two-tailed. P-value of ≤0.05 was considered statistically significant. Correlation analyses were based on linear regression with Pearson correlation. Bar graphs represent mean and standard deviation (SD) unless otherwise stated. All experiments represent at least 3 separate biological repeats, unless otherwise stated.

### Data access

All raw and processed sequencing data generated in this study have been submitted to the NCBI Gene Expression Omnibus (GEO; http://www.ncbi.nlm.nih.gov/geo/) under accession number GSE128999.

## Supporting information

Shani_Erez_Supplementary_Figures

Shani_Erez_Supplementary Table 1

Shani_Erez_Supplementary Table 2

Shani_Erez_Supplementary Table 3

Shani_Erez_Supplementary Table 4

## Acknowledgments

This research was supported by grants to NE from the European Research Council (ERC) under the European Union’s Horizon 2020 research and innovation programme (grant agreement No. 637069 MetCAF), from the Israel Science Foundation (#1060/18), the Emerson Collective, the Israel Cancer Association (ICA) and the Israel Cancer Research Fund (ICRF Project Grant). IT and OM were supported in part by a research grant from the Breast Cancer Research Foundation (BCRF). The authors would like to thank Dr. Ran Elkon for his help with data analysis.

## Author Contributions

OS, YR conceived and carried out experiments; NC, LM, CA, OL-G participated in experiments, OS analyzed data and generated the figures; OM, DS and HS analyzed data; IT and AM were involved in data analysis and discussions; RS, IB and RSS contributed essential resources; NE designed and supervised the study; OS and NE wrote the manuscript.

## Supplementary Tables

**Supplementary Table 1. Related to Figure 3**. Detailed enrichment results for all comparisons based on selection criteria.

**Supplementary Table 2. Related to Figure 3**. Full GSEA results for all comparisons, FDR<0.05, NES>|2|.

**Supplementary Table 3. Related to Figure 4**. List of terms containing TF enriched in all comparisons.

**Supplementary Table 4. Related to Figure 4**. Full results of TF ranking of all comparisons.

