## Supplementary material for "Evolution of metastases-associated fibroblasts in the lung microenvironment is driven by stage-specific transcriptional plasticity": Shani_Erez_Supplementary_Figures

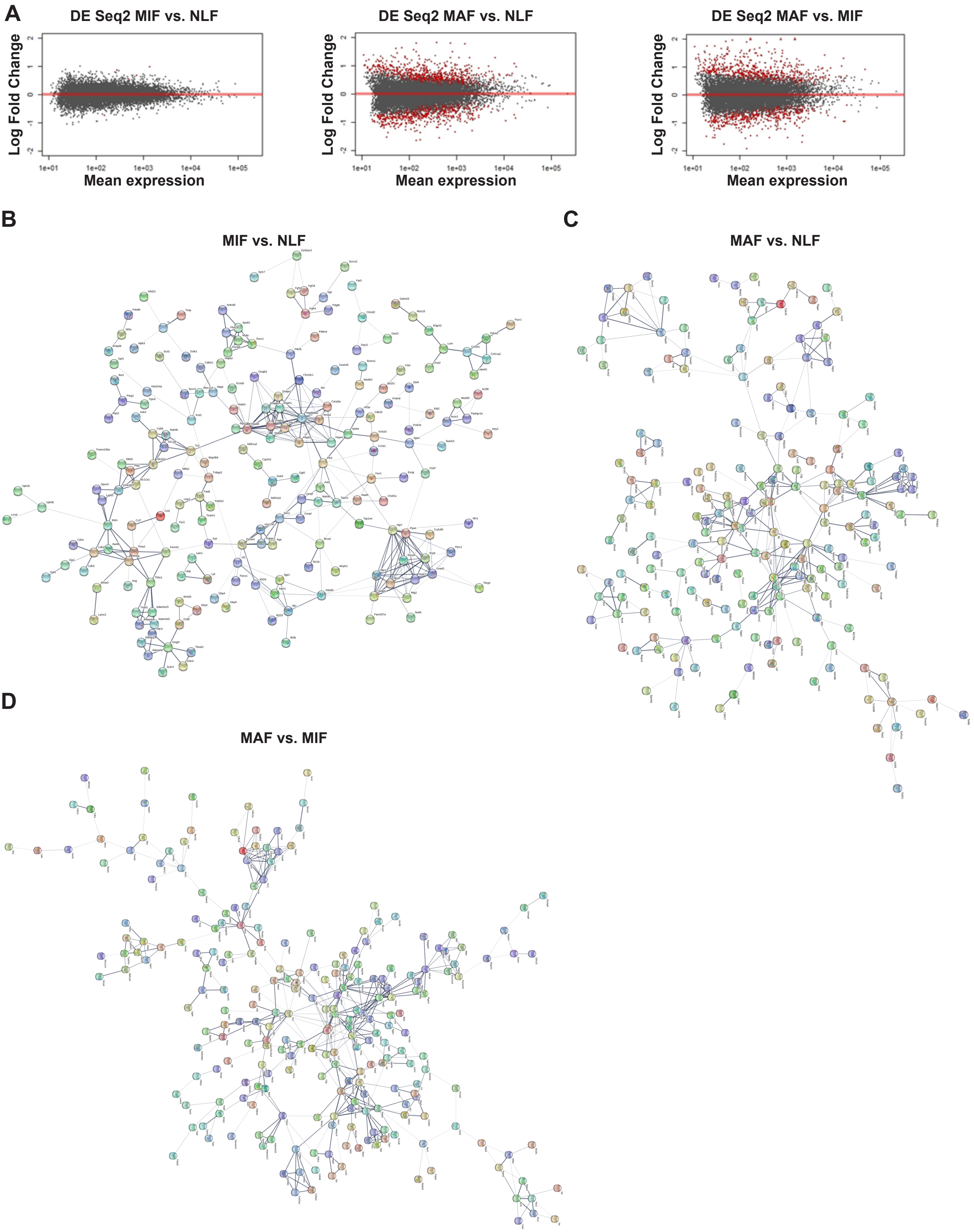


**Supplementary Figure 1. Related to Figure 1. (A)** Volcano plots of differential expression analysis vs. mean expression of MIF vs. NLF, MAF vs. NLF and MAF vs. MIF using DeSeq2. **(B-D)** Protein-protein interactions of differentially expressed genes in each comparison (MIF vs. NLF (B), MAF vs. NLF (C), MAF vs. MIF (D)), derived from the STRING platform. Confidence≥0.3, text mining connections were excluded.

Shani et al. Supplementary Figure 2

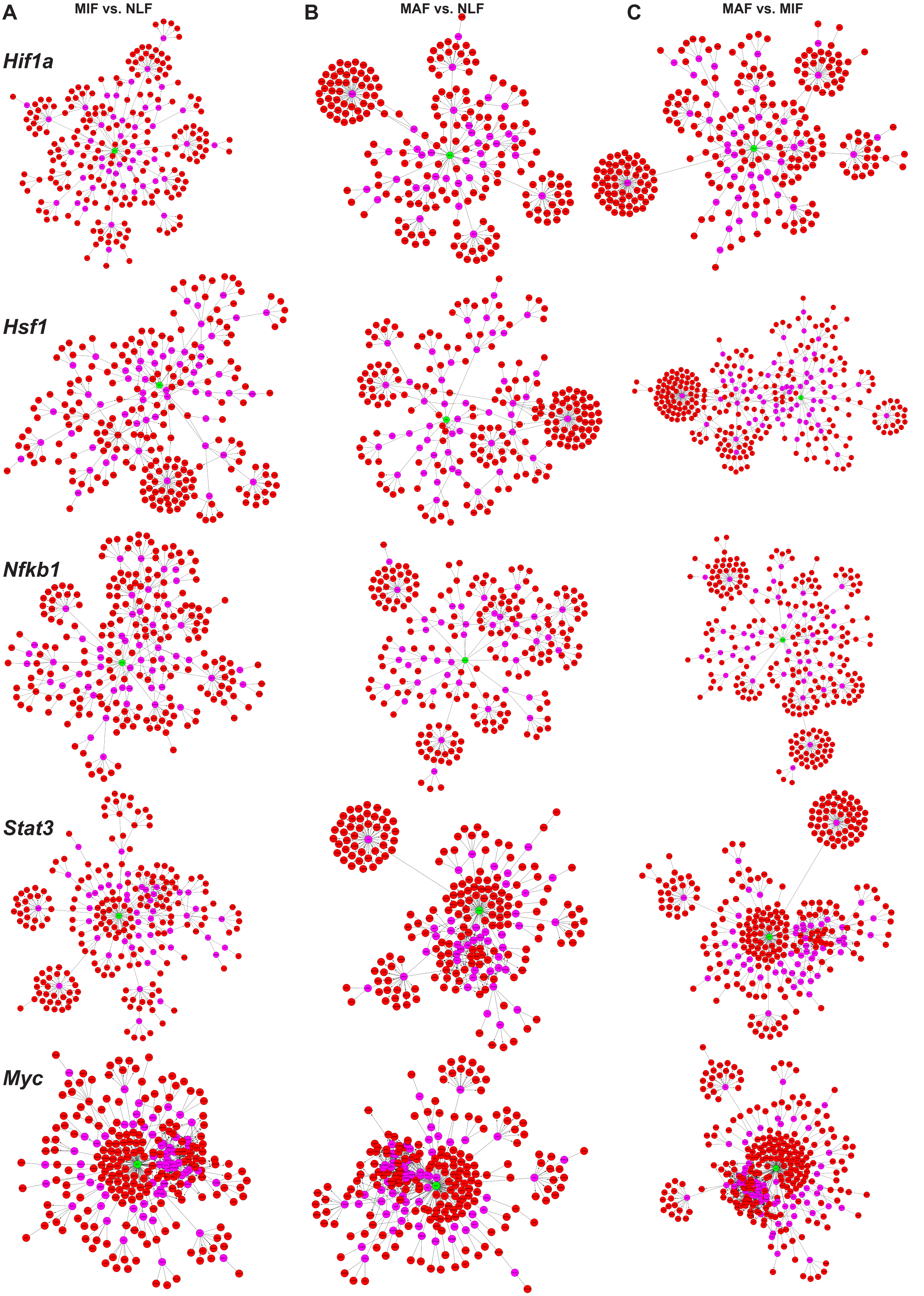


**Supplementary Figure 2**. **Related to Figure 4. (A-C)** ANAT pathway networks for each TF *(Hif1a, Hsf1, Myc, Nfkb1, Stat3)* and each comparison (MIF vs. NLF (A), MAF vs. NLF (B), MAF vs. MIF (C)).

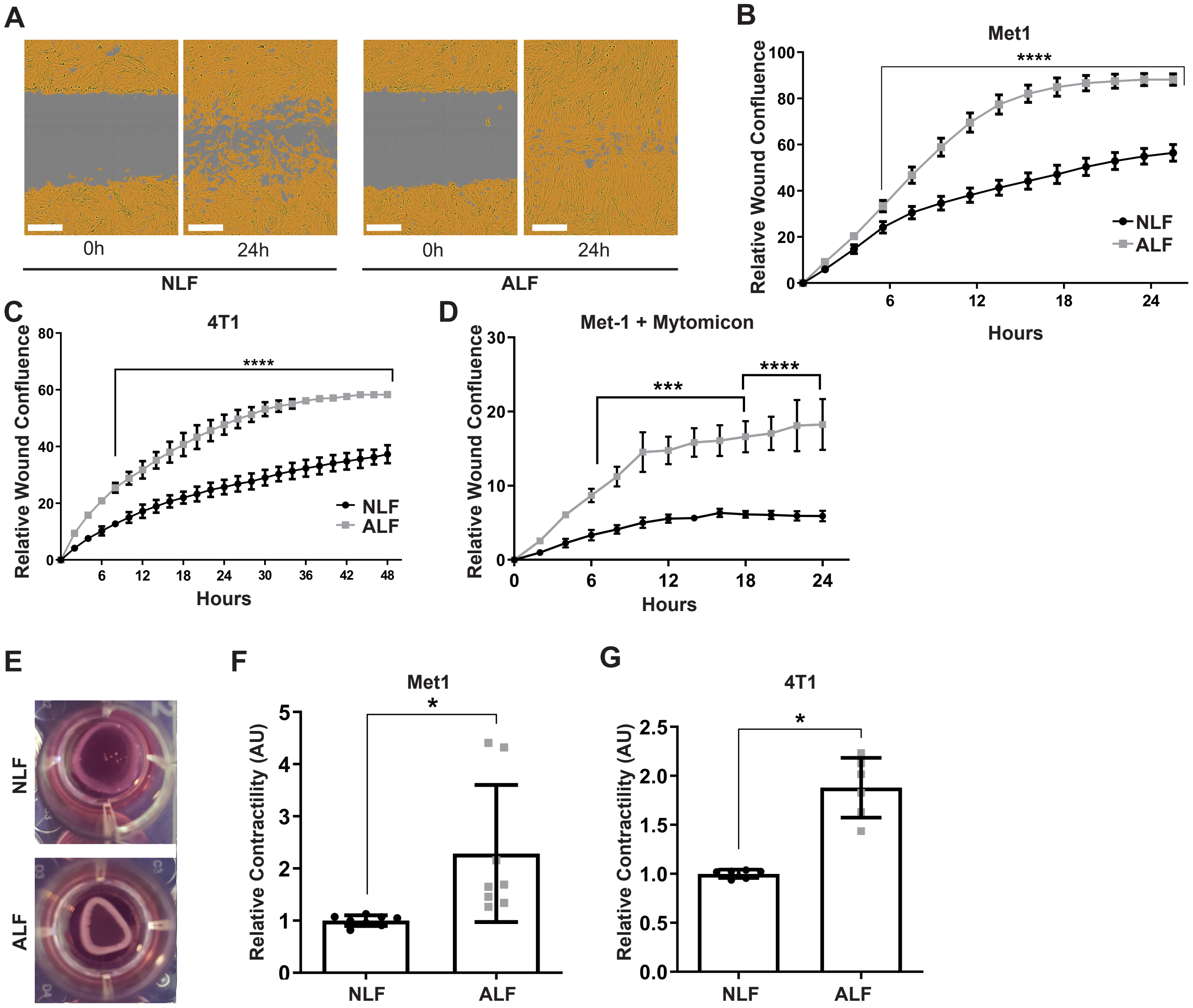


**Supplementary Figure 3**. **Related to Figure 5. (A)** Representative images of scratch closure assay at 0h and 24h following scratch. Lung fibroblasts were incubated with SFM (NLF-normal lung fibroblasts), or with tumor cell CM (ALF-activated lung fibroblasts), scale bar: 300μM. **(B)** Quantification of scratch closure assay performed with FVB/n lung fibroblasts incubated with SFM (NLF, n=3) or with Met1 CM (ALF, n=3) ****$p<0.0001$, Two-way ANOVA with multiple comparisons, data are represented as mean ± SD. **(C)** Quantification of scratch closure assay performed with BALB/c NLF incubated with SFM (n=2) or with 4T1 CM (ALF, n=2), ****$p<0.0001$, Two-way ANOVA multiple comparisons, Data are represented as mean ± SD. **(D)** *Scratch closure is not a result of enhanced fibroblast proliferation:* quantification of scratch closure of lung fibroblasts incubated with SFM (NLF), or with Met1-CM (ALF), and supplemented with the proliferation inhibitor mitomycin C. ***$p<0.001$, ****$p<0.0001$ Two-way ANOVA multiple comparisons, Data are presented as mean ± SEM, n=3. **(E)** Representative images of collagen contraction assay at 24h. Lung fibroblasts were embedded in collagen gel and incubated with SFM (NLF) or in tumor cell CM (ALF). **(F)** Quantification of collagen contraction with FVB/n lung fibroblasts incubated with SFM (NLF, n=8) or with Met1 CM (ALF, n=8), *$p<0.05$, data are represented as mean ± SD. **(G)** Quantification of collagen contraction with BALB/c NLF incubated with SFM (n=2) or with 4T1 CM (ALF, n=2) (E), *$p<0.05$, data are represented as mean ± SD.

Shani et al. Supplementary Figure 4

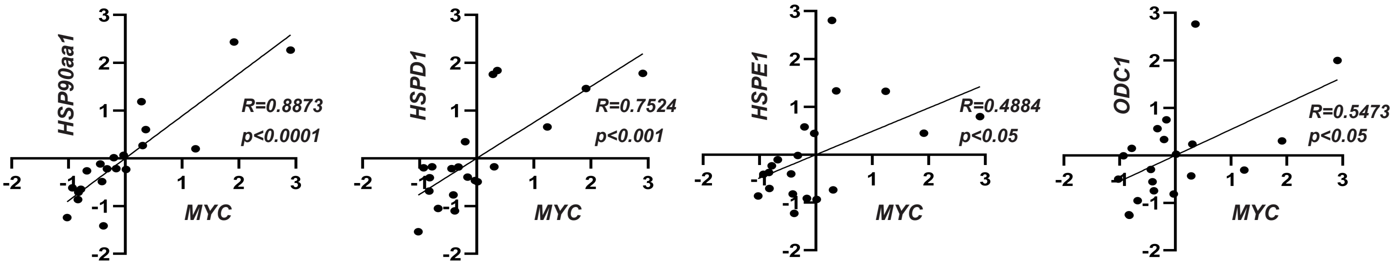


**Supplementary Figure 4, related to Figure 6.** Correlation graphs between *MYC* expression and the expression of specific target genes. P-value of Pearson correlation and correlation coefficient are presented in the graph.
